# From calamity to infestation: linking windstorm tree damage to bark beetle outbreak through forest structure and meteorological analysis

**DOI:** 10.1101/2024.12.05.626970

**Authors:** Michele Torresani, Roberto Tognetti

## Abstract

In recent years, we have witnessed worldwide, an increase in natural forest disturbances, particularly windstorms, which have caused significant direct and indirect forest damages, often triggering largescale bark beetle outbreaks. In this study, we investigated the interaction between windstorm-induced tree damage and subsequent bark beetle outbreaks in the northeastern Italian Alps (Province of Belluno and Bolzano), focusing on the 2018 Vaia windstorm and the successive bark beetle infestation started in 2021. Additionally, we aimed to determine whether this potential correlation is influenced by forest structural characteristics such as forest height heterogeneity (HH), forest density, and forest mean height using LiDAR data, or by meteorological factors (mean temperature and cumulative precipitation) through in-situ spatialized information.

Our research findings, based on a methodology centered on spatial interactions, indicate a potential link between the bark beetle outbreaks and the windstorm event Vaia occurred three years before. Our results suggest that forest structural variables are, in most of the cases, significantly similar across all areas affected by the bark beetle. This similarity is observed both in forests impacted by the Vaia windstorm and in other *Picea abies* forests not affected by the windstorm, indicating that these forest structural variables may not be a trigger for the bark beetle outbreak. Our findings do not show a clear and consistently significant difference in meteorological conditions. This variability can be attributed to the specific areas affected by the Vaia windstorm, which are predominantly mountainous regions characterized by distinct temperatures and precipitation compared to the rest of the provinces. When analyzing the combined influence of structural and meteorological variables in both study areas, our results indicate that none of these factors were ultimately significant predictors of the interaction between bark beetle infestations and areas affected by the Vaia windstorm. Our study suggests that, as climate change increases the frequency and severity of these disturbances, adaptable forest management framework to enhance forest resilience and sustainability are needed, helping forests to better withstand and recover from future natural disturbances.

## 1 Introduction

Over the past few decades, there has been a notable increase worldwide in the frequency and severity of natural forest disturbances [81], such as windthrow [21], drought [60], insect outbreaks [10], and forest fires [14]. These changes, primarily driven by climate change, have significantly affected various forest functions, including ecological processes [82], hydrogeological functions and water regulation [100, 88], forest productivity and carbon sequestration [80], nutrient cycling [48], recreational and aesthetic values [77], and overall biodiversity [101]. These impacts have increased interest in shifting towards smarter forest management practices to enhance the resilience and sustainability of forest ecosystems [90, 66]. Although natural disturbances have historically shaped forest ecosystems, changing disturbance regimes now present an unprecedented threat to vulnerable forests, such as those of mountain environments [8]. The increasing frequency and intensity of these disturbances likely exceed the historic norms of alpine forest ecosystems [5], raising concerns about the sustainability and future of these forests [65].

Bark beetle outbreaks (*Ips typographus*) strongly increased in the northern hemisphere conifer forests in the last four decades mainly due to climate change [31, 80, 44, 40] and, according to available projections, this natural forest disturbance is going to increase in the near future [47, 31]. It has been estimated that this insect is responsible of around 8% of the tree mortality of Europe over the last century [31]. In line with the predisposition and trigger theory [45], bark beetle outbreaks occur when there is a spatial and temporal alignment of vulnerable forests (i.e., those predisposed to stress) with disturbance events that impact tree survival [19]. These stresses include hot and dry weather conditions [47, 56] or windstorm events that cause trees to fall or break and often remain on the ground for extended periods [31, 7]. The fallen trees create an ideal breeding environment for bark beetles, facilitating population explosions and subsequent infestations in surrounding forested areas.

In late autumn of 2018, Italy faced the formidable Vaia windstorm, an atmospheric tempest of unparalleled intensity [98]. Vaia windstorm emerged as a rapid and powerful low-pressure system, producing wind speeds that exceeded typical meteorological expectations. Originating over the Adriatic Sea, the windstorm swept through northeastern Italy, Austria, and Slovenia, carving a path of destruction. With peak wind speeds over 180 km/h [24], Vaia windstorm caused extensive infrastructure damage and severely impacted forested ecosystems, devastating more than 8.5 million cubic meters of wood and largely destroying the mountain ecosystems in its path [93]. Following Vaia, there was a massive amount of fallen trees on the ground; forest management efforts were made to collect the wood as quickly as possible, but in some areas, particularly those difficult to access due to the mountainous terrain, and due to the absence of forest roads, the trees remained on the ground for several years [16]. In some isolated areas, even after six years, the wood still lies on the ground. Physiologically weakened or mechanically damaged trees, whether broken or uprooted, provided ideal breeding grounds for the bark beetle, resulting in a significant increase in beetle populations [16] that started in 2021. This temporal pattern has been observed in other related studies, where beetle populations typically peaked in the third growing season following a disturbance [16, 57, 81]. Different studies have highlighted the clear connections between wind-induced forest damages and subsequent bark beetle outbreaks [25, 58, 32]. In the case of Vaia windstorm, some studies have focused on various linked aspects, such as the map spatialization of the bark beetle outbreak between 2015 and 2021 in a restricted area of the Belluno area [54], and the semiochemical push-and-pull technique [16] for reducing bark beetle damage in Vaia-affected areas. However, the potential link between the Vaia windstorm and the bark beetle outbreak, including the influence of forest structural variables (forest height heterogeneity – HH -, forest density and forest mean height) and meteorological factors (such as temperature and precipitation), has not been thoroughly analyzed.

The advent of remote sensing technologies, including optical, LiDAR, and radar sensors, alongside advancements in platforms such as airborne, satellite, and UAV systems, has revolutionized our ability to detect and extract crucial information for ecosystem monitoring [91, 95, 75, 36, 11, 76, 9], especially regarding natural forest disturbances [64, 83, 78, 42, 37, 3, 27, 86]. The efficiency of these technologies is evident in their ability to swiftly cover vast areas, providing insights that would be logistically challenging and resource-intensive to obtain through traditional on-the-ground surveys alone [91, 63]. However, it is important to note that while remote sensing offers speed and cost-effectiveness, there can be a trade-off in terms of reduced accuracy compared to meticulous field data collection [71]. Despite this, numerous studies have demonstrated the utility of remote sensing data in monitoring various forest disturbances, such as wind damage [103, 30], forest fires [15, 4, 85], droughts [51, 105, 104], and infestations [39, 2, 42, 22], underscoring its importance in ecosystem management. LiDAR data, in particular, enhances our understanding of forest ecosystems by providing detailed information on structural attributes, biodiversity metrics, topographical features, forest biomass, canopy height, and vegetation density [67, 6, 18, 26]. This technology is instrumental in forest management, offering a rapid and cost-effective alternative to traditional field-based methods. Additionally, LiDAR’s capability to capture detailed three-dimensional information enables precise quantification of forest structure and assists in monitoring changes over time [94].

The primary aim of this study is to investigate the potential interaction between the damage caused by the Vaia windstorm, which occurred in 2018, and the subsequent bark beetle outbreak of 2021 in northeastern Italy, particularly in the Province of Belluno (Veneto region) and in the Province of Bolzano (South Tyrol region). In addition we aim to understand if this potential correlation is influenced by structural characteristics of the forest (forest HH, forest density and forest mean height) through the use of LiDAR data, or by meteorological factors using in-situ spatialized information. Information on forest structure and status before and after the Vaia windstorm and beetle outbreak will help understand trends over time and implement management approaches to reduce the current vulnerability of mountain forests to these disturbances [12, 46].

## 2 Materials and methods

### 2.1 Study areas

Our approach was tested in Italy, in northeastern Italy, particularly in the Province of Belluno (Veneto region), and in the Province of Bolzano (South Tyrol region). The Province of Belluno covers an area of 3,672 square kilometers, it extends from the Dolomite Alps in the north to the Venetian Prealps in the south. The elevation varies significantly, ranging from 42 meters to 3,325 meters above mean sea level. Forest covers around 60% of the area. Province of Bolzano is located at the west side of Belluno, it encompasses an area of approximately 7,400 km², with elevations ranging from 194 meters to 3,905 meters above sea level. As the area of Belluno, Bolzano is predominantly mountainous and widely covered by forests (50% of the total area). The choice to analyze data from these two areas is due to the unique position as the only Italian regions that provides open access to data on bark beetle infestations over time.

Both areas were heavily hit by the Vaia windstorm; according to the Forwind database [23] (see sub-section 2.3.2 Forwind data-set) the forest area lost by the area of Belluno was approximately 2700 hectares. The windstorm predominantly impacted the northwestern part of the Province, particularly the Agordo area, as well as some regions in the northeast, while other areas remained unaffected (Figure 2). The area of Bolzano lost approximately 6,000 hectares of forest. Vaia windstorm hit mainly the western part of the Province. This significant loss of forest cover not only disrupted the local ecosystem but also increased the vulnerability of the remaining trees to subsequent disturbances, such as bark beetle infestations (Figure 1).

**Figure 1:**
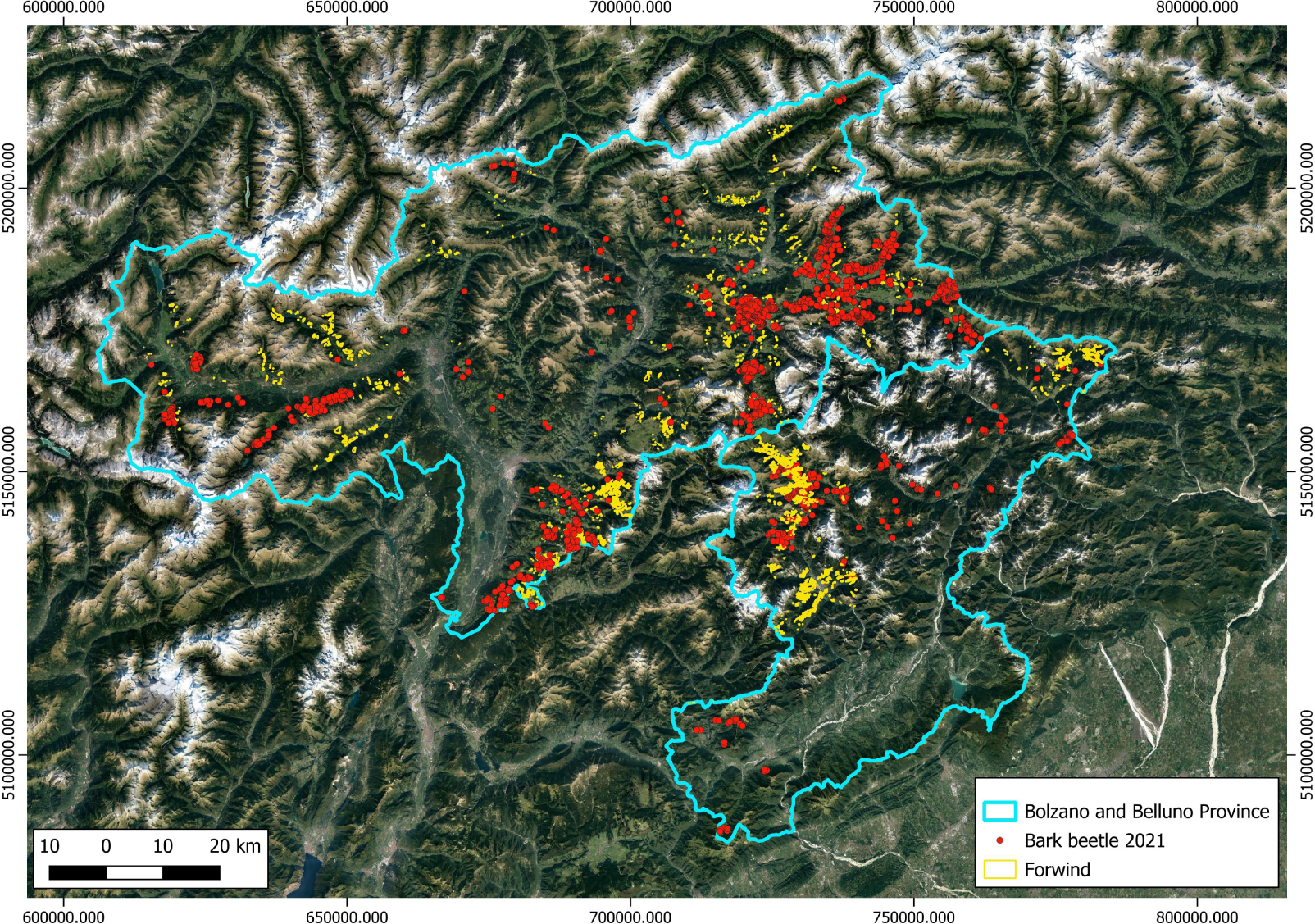
In blue the border of the two study areas. On the east, the Belluno Province and on the west, the Bolzano Province. In yellow the forest hit by the Vaia windstorm as identified by the Forwind dataset and in red the point location of bark beetle infestation of 2021, identified by the local forest services.

**Figure 2:**
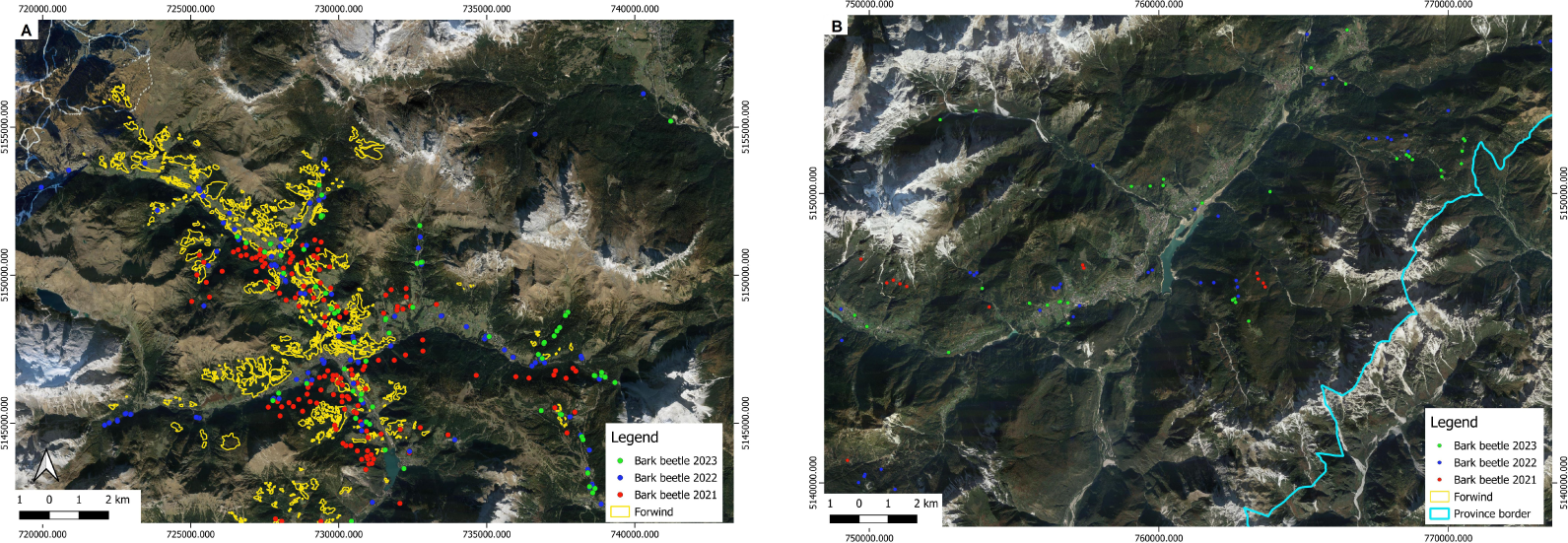
An illustration of the varied impact of the Vaia windstorm and subsequent bark beetle infestation within the same area (in this case Belluno area). Sub-plot A (Agordo area, in the North-west side of the area) shows an area heavily hit by the Vaia windstorm in 2018 and successively by the bark beetle outbreak. Sub-plot B (Cadore area, in the North-west side of the area) shows an area not damaged by Vaia windstorm and not heavily hit by the bark beetle outbreak.

### 2.2 Workflow and statistical analysis

#### 2.2.1 Assessment of possible interaction between bark beetle affected areas and forest stands hit by Vaia windstorm

The methodology involved several steps to ensure robust statistical analysis and minimize possible bias (Figure 3). First, for each study areas (Belluno and Bolzano), we selected 500 random points within the areas affected by the Vaia windstorm, as identified in the Forwind dataset (see sub-section 2.3.2 Forwind data-set) [23]. For each of these points, we calculated the altitude and categorized them into three classes: 500-1100 m, 1100-1700 m, and 1700-2300 m above mean sea level (a.m.s.l.). Subsequently, a buffer of 500 meters was created around each of these points. Next, we selected 500 random points in *Picea abies* forests (identified using the vector layer “Carta Regionale dei tipi forestali” for the Belluno area, and through the vector layer “Tipologie forestali” for the Bolzano area, which are both freely available online) not affected by the Vaia windstorm. The selection of these points was stratified to match the altitude classes of the Vaia windstorm points. Specifically, for each point in the Vaia Forwind dataset, we selected a corresponding random point within the same altitude class in the unaffected forests. This stratified random sampling was employed to eliminate altitude-related bias. A 500 m buffer was also created around each of these points in the unaffected forests. We then conducted an intersection analysis with the bark beetle infestations recorded in 2021 (see sub-section 2.3.1 Bark beetle data infestation). This analysis was performed to identify possible intersections between the bark beetle infestations (punctual information) and the 500 m buffers of both random points within the Vaia windstorm-affected areas and the random points in other *Picea abies* forests unaffected by Vaia. Subsequently, we checked the number of points (500 for Vaia and 500 for *Picea abies* non-Vaia) that had at least one intersection with the bark beetle infestations. We determined which variable (Vaia or *Picea abies* non-Vaia) had more interactions (more than 0) and performed a t-test to assess the significance of the difference. This analysis was repeated 100 times, each time selecting 1000 new random points (500 for Vaia and 500 for *Picea abies* non-Vaia) to ensure a robust statistical analysis.

**Figure 3:**
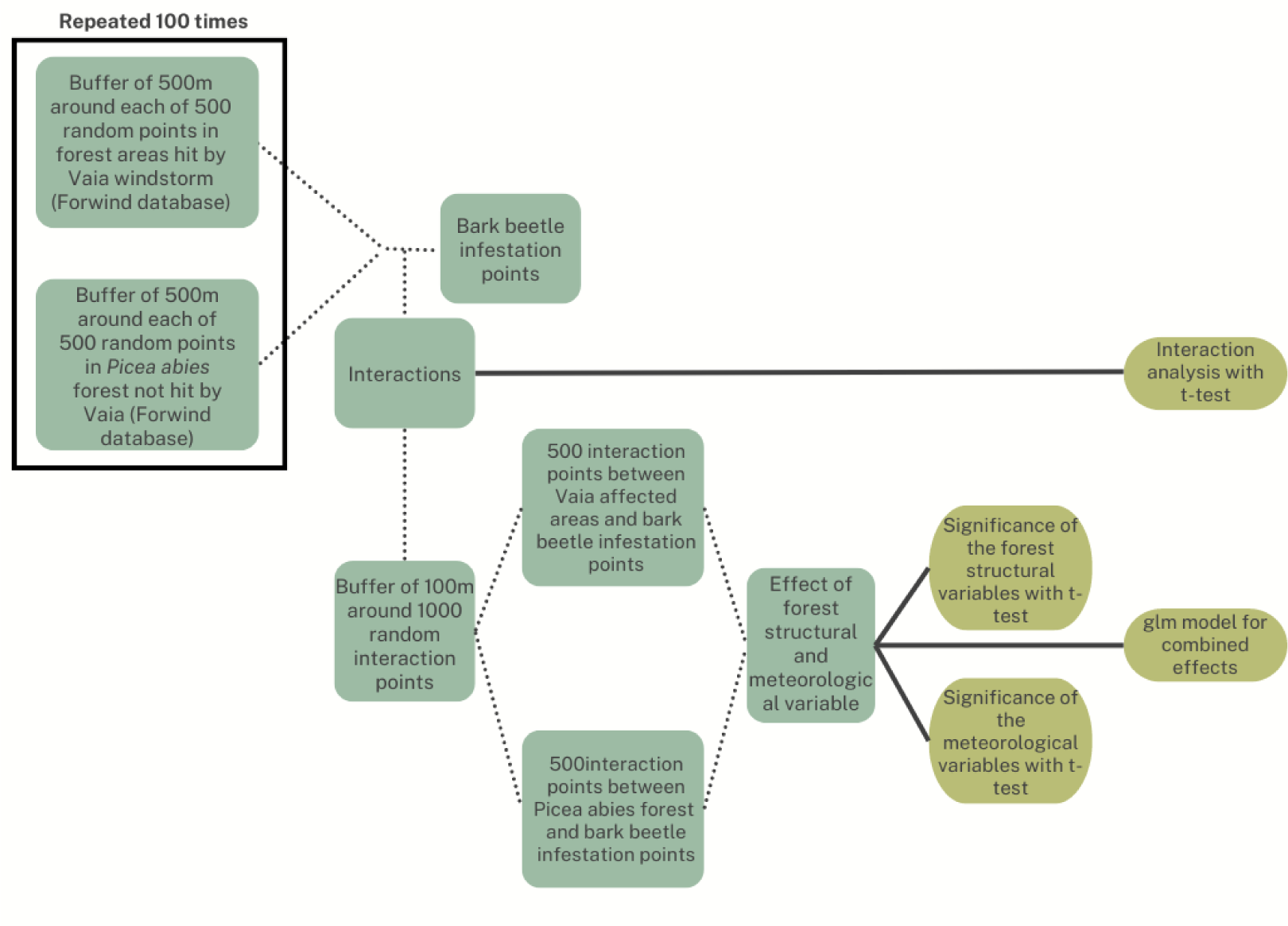
The proposed approach.

#### 2.2.2 Assessment of the role of forest structural and meteorological variables on the possible interaction between bark beetle affected areas and forest stands hit by Vaia windstorm

In both selected areas of study, for all positive intersections between the bark beetle infestations and both areas affected by the Vaia windstorm and the *Picea abies* forests not affected by Vaia (derived from the 100 random repetitions), we selected an additional 1000 random points (500 within the intersections with Vaia windstorm-affected areas and 500 within the intersections with *Picea abies* forests not affected by Vaia). For each of these points, a virtual buffer of 100 meters was created. For each of the 1000 buffers areas (having g 100m of buffer), we derived three different forest structural variables (forest HH, forest density and forest mean height) estimated through Canopy Height Model (CHM - based on airborne LiDAR information - see sub-section 2.3.3 LiDAR Data and sub-section 2.4 Forest Structural Variables). For each point we also derived (at a scale of 1km) the mean values of mean temperature and cumulative precipitation (see sub-section 2.3.4 Meteorological data). We compared the forest structural and meteorological variables between the infested bark beetle areas hit by Vaia windstorm and the infested bark beetle areas in *Picea abies* forests not affected by Vaia windstorm. We performed a t-test for each distinct variable, to assess the significance of the differences, aiming to determine whether these variables could have influenced the relationship between the Vaia windstorm and the subsequent bark beetle outbreak.

Finally we performed a logistic regression analysis (glm model) to investigate the combined effects of forest structural variables and meteorological variables. The analysis was done for both study areas, including the 1000 random points for each study area (2000 in total), derived from positive intersections of bark beetle infestations and respective forest areas.

### 2.3 Used data

#### 2.3.1 Bark beetle data infestation

In the area of Belluno, the bark beetle infestation data were downloaded from the open access geo portal “Geoportale dei dati territoriali” (https://idt2.regione.veneto.it/idt/webgis/viewer?webgisId=204). For the spatial interaction analysis we used the punctual information, collected on the ground, of the “Bark beetle infestation site” of 2021, which was the earliest available temporal dataset. To obtain a broader estimate of the areas affected by the bark beetle over time, we also accessed data from the “monitoring of probable areas affected by bark beetle” for 2021 and 2022 (the only available online), which was developed and provided by the Veneto region [99].

In the area of Bolzano, the bark beetle infestation data were downloaded from the open access geo portal “Geocatalogo” (http://geokatalog.buergernetz.bz.it/geokatalog/#!) using the “Forest damage caused by the spruce bark beetle” (Bostrico dell’abete rosso, Ips typographus) vector layer, for the years 2021 and 2022. These data were utilized as point information for interaction analysis (using the centroid of each available polygon) and for estimating the areas affected by the bark beetle over time.

#### 2.3.2 Forwind data-set

The Forwind data-set [23], which is freely available here (https://figshare.com/articles/dataset/A_spatially-explicit_database_of_wind_disturbances_in_European_forests_over_the_period_2000-2018/9555008), was utilized in this study to characterize the area affected by the Vaia windstorm (Figures 1). This dataset provides a novel repository documenting wind disturbances in European forests, encompassing over 80,000 spatially delineated areas that experienced wind-related disturbances between 2000 and 2018 [23]. Presented in a standardized geographical vector format, the dataset ensures consistency and comprehensiveness. It includes all significant windstorms recorded during the observational period, with the Vaia windstorm accounting for approximately 30% of reported damaging wind events in Europe. To validate the reliability of the Forwind data, correlation analyses with land cover changes derived from the Landsat-based Global Forest Change dataset and the MODIS Global Disturbance Index were conducted, confirming the robustness and accuracy of the Forwind dataset [23].

#### 2.3.3 LiDAR data

In order to estimate the effect of forest structural variables on the possible correlation between the Vaia windstorm and the subsequent bark beetle outbreak we made use of an Airborne Laser Scanner (ALS) LiDAR dataset conducted in both areas prior to both forest disturbances.

In the area of Belluno, we used the CHM (available online by the area of Belluno here https://www.regione.veneto.it/web/agricoltura-e-foreste/ modelli-digitali-delle-chiome) with a spatial resolution of 1m, derived from an ALS LiDAR campaign carried out in 2015.

In the area of Bolzano we used the ALS LiDAR data derived from a campaign conducted in 2006; we derived the CHM (with a spatial resolution of 2.5m, the most accurate available) as the difference between the DSM and DTM both available online in the open access geo portal “Geocatalogo” (http://geokatalog.buergernetz.bz.it/geokatalog/#!).

#### 2.3.4 Meteorological data

For the area of Belluno, the spatialized cumulative precipitation and average temperature data were provided by ARPA Veneto (Regional Agency for Environmental Protection). The data, covering the period from 2018 to 2021, have a spatial resolution of 1 km. These data were derived from the spatial interpolation of all available meteorological stations using the Universal Kriging method [28].

For the area of Bolzano, the data, with the same spatial resolution (1km), covering the same period (2018-2021) were provided by the EURAC research center. Also in this case the data were derived from the spatial interpolation of all available meteorological stations using the methodology described by Crespi et al. [13].

### 2.4 Forest structural variables

To estimate the effect of forest structural variables on the possible correlation between the Vaia windstorm and the subsequent bark beetle outbreak, we determined whether the forest structure, including forest HH, forest density, and forest mean height (Table 1) in areas where bark beetle infestation intersected with Vaia windstorm-affected regions was similar to or different from that in bark beetle-infested areas not near Vaia windstorm-affected zones. If the structures were significantly similar, it would suggest that the triggering effect could have been the Vaia windstorm itself. Conversely, if the structures were significantly different, it would indicate that the forest structure might be a triggering variable of the outbreak. For this analysis, in both studied areas, we selected 500 random points from the intersections between Vaia and bark beetle infestation areas and another 500 random points from the *Picea abies* forests not affected by Vaia but infested by bark beetles. This selection considered the 100 iterations of the intersection analysis. For each of these points, we calculated the three forest structural variables within a virtual 100 m buffer (1ha each), using LiDAR data.

**Table 1:** The table summarizes the structural variables used in this study (Forest Height Heterogeneity - HH -, forest density and forest mean height).

| Structural Variable | Explanation |
| --- | --- |
| Forest height heterogeneity (HH) | Calculated as the heterogeneity (through the Rao's Q index) of CHM pixel values |
| Forest density | Calculated through the canopy cover, as the ratio between the number of CHM pixels >2m and the total number of pixels |
| Forest mean height | Calculated as the mean CHM values |

#### 2.4.1 Forest height heterogeneity

HH expresses the diversity in forest tree height derived from CHM LiDAR data [96]. This measure, based in the Height Variation Hypothesis approach (analogous to the Spectral Variation Hypothesis that uses optical data [72]), states that high HH values correlates with habitat having higher environmental heterogeneity and biodiversity. As demonstrated in various studies [96, 94, 87], HH serves as an indicator of forest vertical structure, intricately linked to habitat diversity and biodiversity [52, 53]. The computation of HH need the use of an heterogeneity index, which effectively measures the diversity within CHM pixel values. In this context, the HH in areas where bark beetle infestation intersected with Vaia-affected regions and in bark beetle infested areas not near Vaia-affected zone was derived, for its proven effectiveness [72, 69, 70, 68], using the Rao’s Q index (Eq.1). The suitability of this index for calculating HH using CHM had been previously validated with robust results in other studies [50, 87].

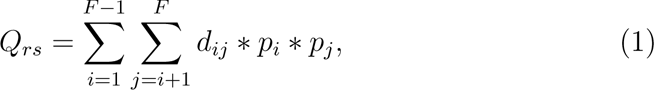

where:

*Q_rs_* = Rao’s Q index

*p* = relative abundance of a CHM pixel value in a given area (F)

*d_ij_* = distance between the i-th and j-th pixel value (*d_ij_* = *d_ji_* and *d_ii_* = 0)

i = pixel value i

j = pixel value j

The relative abundance *p* is calculated as the ratio between the considered pixel value (*p_i_* and *p_j_*) and the total number of pixels in F. As well discussed by Rocchini et al. [72], the index allows for the construction of a distance matrix *d_ij_* in different dimensions, enabling the consideration of more than one band or raster at a time. In our case, and as used in different other studies [96, 87], the *d_ij_* was calculated as a simple Euclidean distance based on a single layer (CHM raster values).

In both studied areas, the Rao’s Q values were retrieved for each 100 m buffer (1ha each) area of the new random points (500 from the intersections between Vaia windstorm and bark beetle infestation and another 500 random points from the intersection between the *Picea abies* forests not affected by Vaia windstorm but infested by bark beetles.) implementing the R spectralrao function (of the rasterdiv R package [74]).

#### 2.4.2 Forest density

Following the results of previous works [96, 94], the forest density was calculated in both areas, as canopy cover for both groups of the 500 new random areas (100 m of buffer) through the following formula:

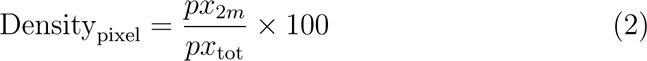

where:

*Density_pixel_*= forest density

px_2*m*_= number of pixels with a CHM > 2m

px*_tot_*= total number of pixels

#### 2.4.3 Forest mean height

In both studied areas, for both groups of the 500 new random areas (100 m of buffer), the forest mean height was computed by calculating the mean CHM values (Eq. 3).

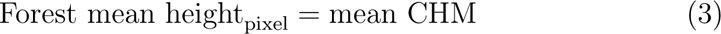

where:

Forest mean height_pixel_= forest mean height

mean CHM= mean of CHM (Canopy Height Model) values per plot

## 3 Results

Figure 4 shows, for both study areas, how many times out of 100 iterations the 500 m buffer zones around the 500 points within the Vaia windstorm-affected areas (as identified in the Forwind dataset) had more interactions with bark beetle infestations, compared to the other 500 m buffer areas around the 500 random points within other *Picea abies* areas not affected by Vaia windstorm.

**Figure 4:**
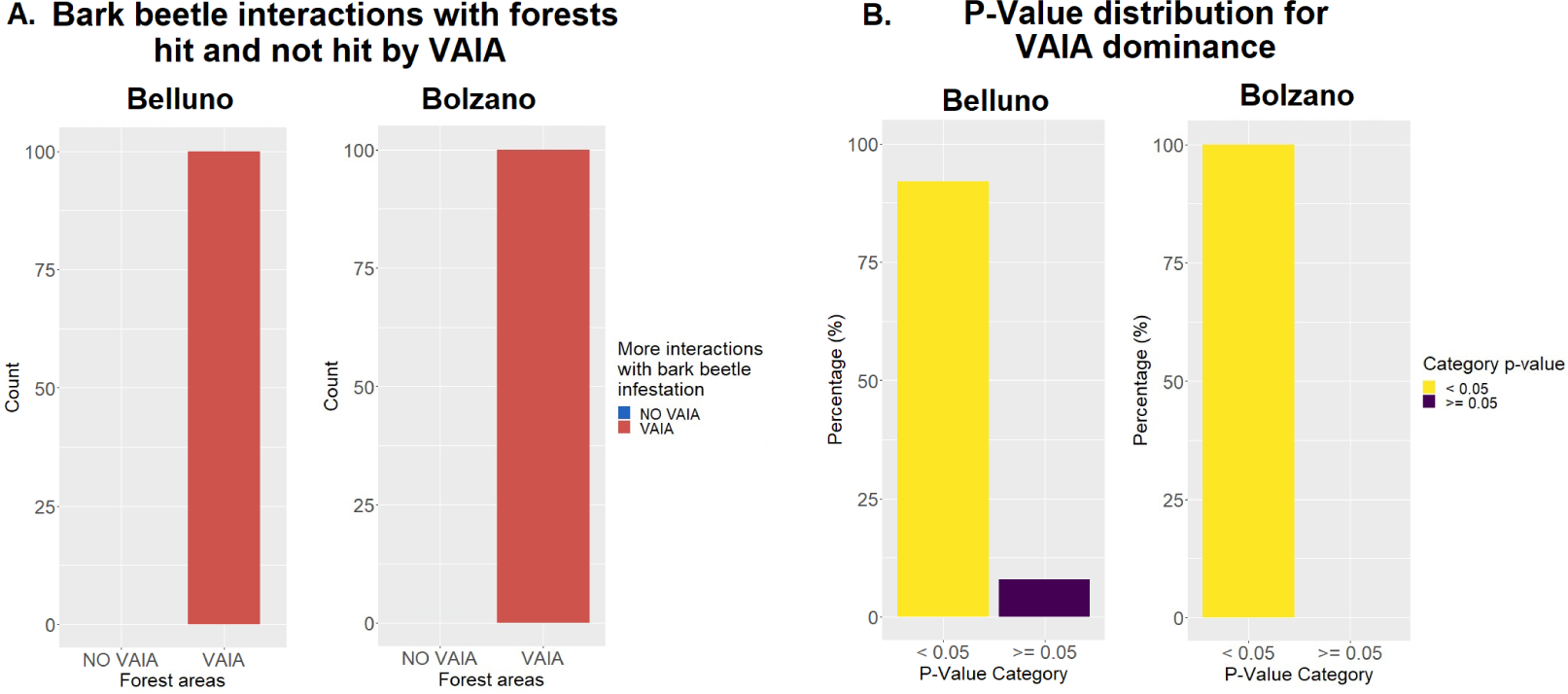
Sub-plot A illustrates, for both study areas, the number of iterations (out of 100) where there was a higher interaction between the areas affected by bark beetle in 2021 and 500 random points within the Vaia windstorm-affected areas (VAIA) or 500 random points within other *Picea abies* forested areas (NO VAIA). Sub-plot B displays the significant percentage of iterations (for the VAIA areas) corresponding to sub-plot A.

The results indicate that, in both study areas, in 100% of the iterations, the bark beetle infestation areas, showed more interactions with the Vaia windstorm-affected areas than in the other *Picea abies* forests not affected by Vaia windstorm.

Focusing on the Vaia iterations, nearly 85% of iterations for Belluno and 100% for Bolzano were statistically significant.

Figure 5 shows, the differences in mean temperature and cumulative precipitation in both study areas. These measurements were taken within each 500 random points where interactions between the bark beetle infestation and the Vaia windstorm-affected areas occurred, and within the 500 random points where interactions between the bark beetle infestation and other *Picea abies* forests occurred. This analysis was conducted to determine if these meteorological variables could be considered additional triggers for the bark beetle infestations. The results show, for both study areas that the mean temperature is statistically similar (p-value of t-test >0.05) underlying that, it can not be considered an additional trigger for the bark beetle infestation. The cumulative precipitation was significantly different in both the areas. It was higher in the Vaia affected areas in the Province of Bolzano while lower in the Province of Belluno.

**Figure 5:**
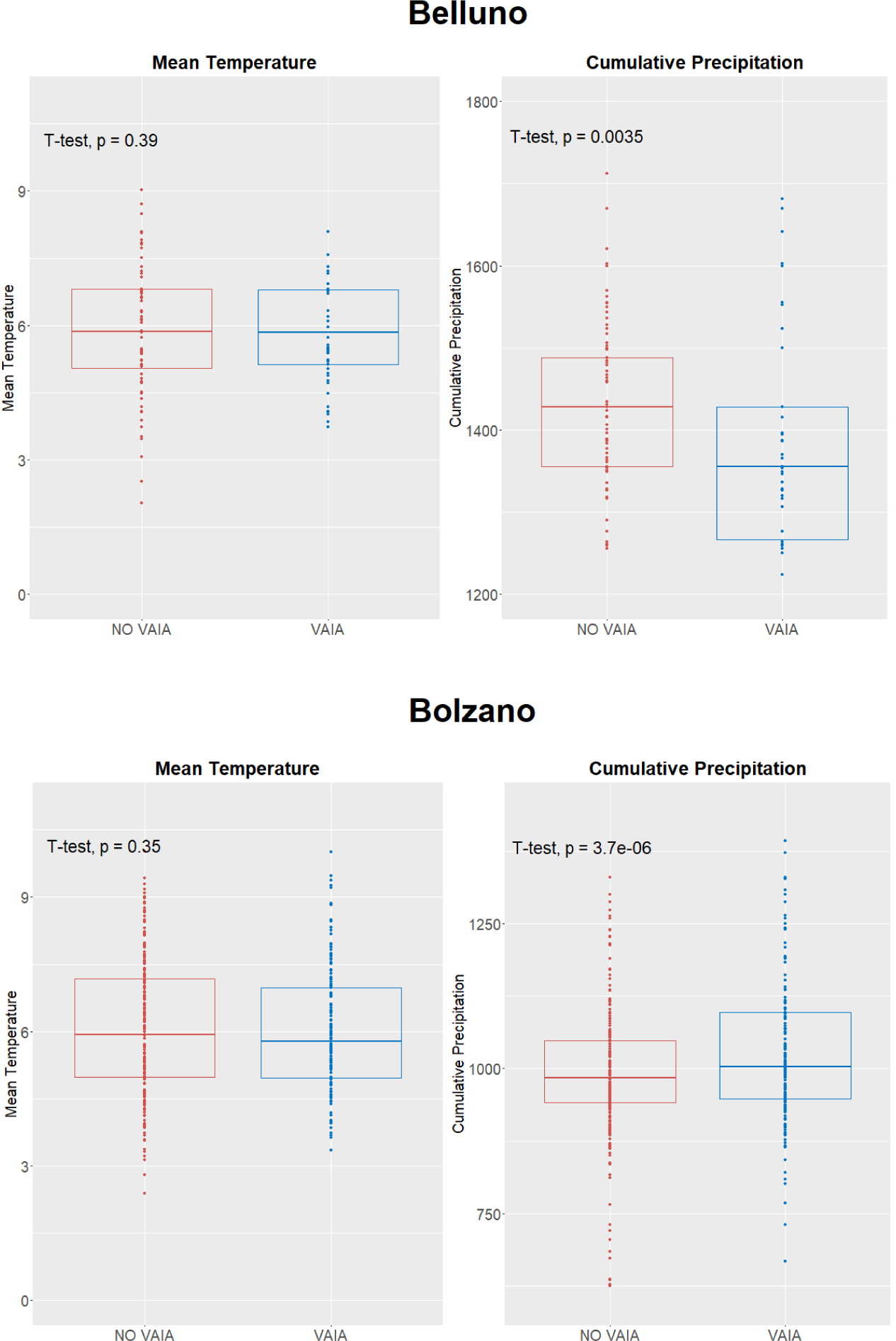
Differences in mean temperature and cumulative precipitation for the 500 random points where interactions between bark beetle infestations and Vaia windstorm-affected areas occurred, and for the 500 random points where interactions between bark beetle infestations and other *Picea abies* forests occurred. The analysis shows that the mean temperature is statistically similar in both areas (p-value of t-test > 0.05), indicating that this meteorological variable may not be an additional trigger for the bark beetle infestations. Conversely, cumulative precipitation shows a significant difference in both areas.

Figure 6 shows, for both study areas, the differences in forest HH, forest density and forest mean height calculated with LiDAR CHM data. These measurements were taken within the 100 m buffer (1ha each) zones around the 500 random points where interactions between the bark beetle infestation and the Vaia windstorm-affected areas occurred, and within the 100 m buffer (1ha each) zones around the 500 random points where interactions between the bark beetle infestation and other *Picea abies* forests occurred. This analysis was conducted to determine if these structural variables could be also considered additional triggers for the bark beetle infestations. The results show, for both study areas that forest density and forest mean height are statistically similar (p-value of t-test >0.05) underlying that, these factors should not be considered an additional trigger for the bark beetle infestation. Only the forest HH heterogeneity in the Belluno area showed a significance difference between the two distinct areas.

**Figure 6:**
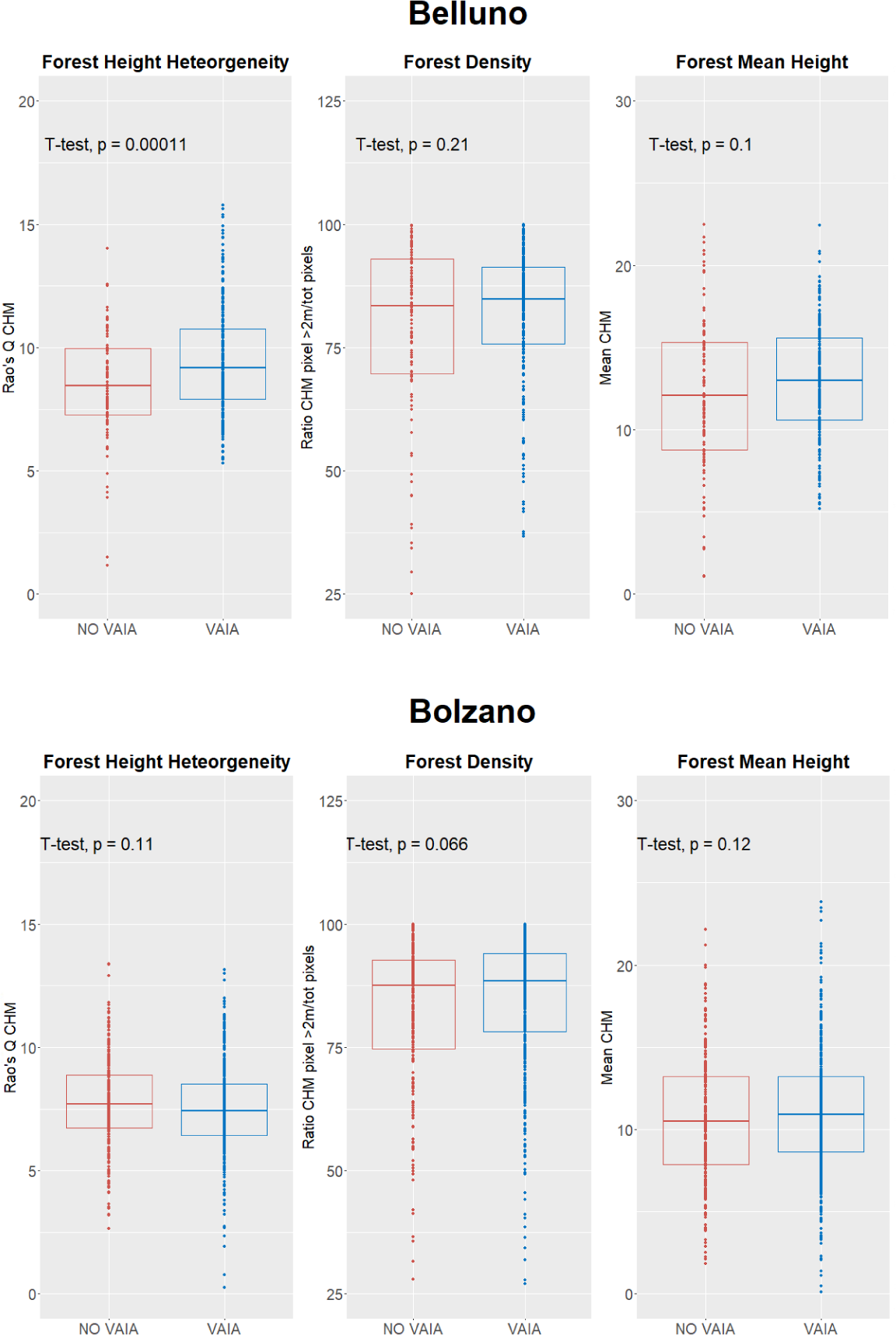
Differences in forest Height Heterogeneity (HH), forest density, and forest mean height calculated using LiDAR CHM data within the 100 m buffer (1ha each) zones around 500 random points where interactions between the bark beetle infestation and Vaia windstorm-affected areas occurred, and within the 100 m buffer (1ha each) zones around 500 random points where interactions between the bark beetle infestation and other *Picea abies* forests occurred. The analysis shows for both study areas, that forest density, and forest mean height are statistically similar in both areas (p-value of t-test > 0.05), indicating that these structural characteristics may not be additional triggers for the bark beetle infestations. Only the forest HH showed, for the Belluno area, a significant difference.

Finally a logistic regression analysis (glm model) was performed to investigate the combined effects of forest structural variables and meteorological variables on the likelihood of bark beetle infestation in areas affected by the Vaia windstorm versus *Picea abies* forests not affected by the windstorm. The analysis for both study areas, included the 1000 random points for each study area (2000 in total), derived from positive intersections of bark beetle infestations and respective forest areas. The results are summarized in Table 2. Overall, the model suggests that none of the variables considered show a strong and statistically significant effect on the likelihood of bark beetle infestation when comparing Vaia-affected areas to unaffected *Picea abies* forests.

**Table 2:** Summary of the logistic regression analysis (glm model) assessing, on both study areas, the combined effect of forest structural variables (forest height heterogeneity, forest mean height, and forest density) and meteorological variables (cumulative precipitation and mean temperature) on the likelihood of bark beetle infestation in areas affected by the Vaia windstorm versus *Picea abies* forests not affected by the windstorm.

| Variable | Estimate | Std. Error | z value | Pr(>\|z\|) |
| --- | --- | --- | --- | --- |
| (Intercept) | -0.3008 | 0.7410 | -0.406 | 0.685 |
| Forest HH | 0.0192 | 0.0389 | 0.493 | 0.622 |
| Forest mean height | 0.0050 | 0.0300 | 0.166 | 0.868 |
| Forest density | 0.0101 | 0.0063 | 1.609 | 0.108 |
| Mean precipitation | 0.0004 | 0.0003 | 1.091 | 0.275 |
| Mean temperature | -0.0737 | 0.0535 | -1.378 | 0.168 |

Figure 7 illustrates the forest area (in hectares) for both study areas, that was damaged by the Vaia windstorm in 2018, as recorded in the Forwind dataset and the subsequent area affected by bark beetle infestations in 2021 and 2022, based on freely available data provided by both studied Provinces (see sub-section 2.3.1 Bark beetle data infestation). The graph indicates that the bark beetle outbreak over just two years has caused nearly as much damage as the Vaia windstorm in the Belluno area and even more damage in the Bolzano area.

**Figure 7:**
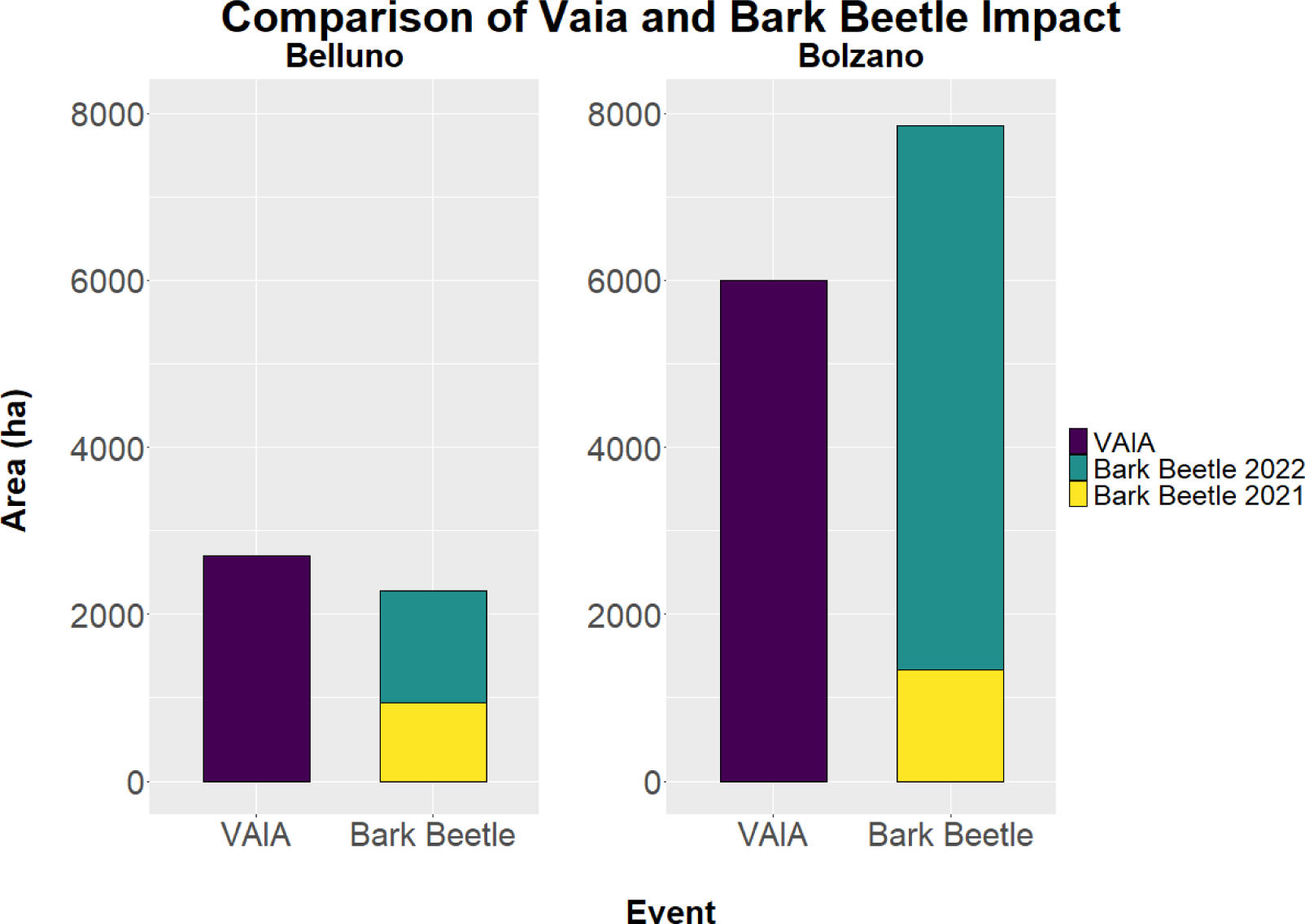
Estimated area (ha) affected by the Vaia windstorm of 2018 (Forwind dataset) and damaged by the bark beetle outbreak of 2021 and 2022 for both studied areas (Belluno and Bolzano).

## 4 Discussion

In this study, we explored the potential correlation between the forest damages caused by the Vaia windstorm, which significantly impacted the northeastern Italian forests in 2018, and the subsequent bark beetle outbreak occurred starting from 2021/2022 in the Provinces of Belluno and Bolzano (northeastern Italian Alps). In addition, we investigated the role of meteorological variables (temperature and precipitation) and forest structural variables (HH, forest mean height and forest density assessed through ALS LiDAR data) on the above mentioned correlation. Our findings, utilizing a methodology centered on spatial interactions, indicate a potential correlation between forest damage caused by the Vaia windstorm and subsequent bark beetle outbreaks in both study areas. This correlation has been extensively documented in various regions affected by similar events. For instance, bark beetle outbreaks were observed following the Kyrill hurricane [102] and the Elisabeth and Phillip storms in the Czech Republic [20], the Alžbeta windstorm in Slovakia [58], and the Gudrun windstorm in Sweden [31]. However, our study marks the first analysis of this correlation in the context of the Vaia windstorm. Windstorms, such as Vaia, cause significant tree damage, creating a large amount of fallen and weakened trees that provide ideal breeding conditions for bark beetles [25]. These insects are highly opportunistic and thrive on stressed or dead wood, which is abundant after a windstorm. The fallen trees serve as a rich resource for bark beetles to reproduce, leading to a rapid increase in their population. As these beetles mature, they expand their range to healthy trees, further exacerbating forest damage [25]. The aftermath of a windstorm, in addition to impacting regeneration processes naturally adapted to small-scale canopy openings [41], significantly influences the extent and severity of beetle outbreaks. The large amount of downed wood creates a highly favorable environment for beetle infestation and proliferation. Furthermore, the legacy of a windstorm can alter resource availability crucial for recovery post-beetle outbreak, constraining a forest’s resilience against subsequent disturbances, including wildfires [46].

As highlighted in various studies [32], tree mortality caused by *I. typographus* following windstorms can increase exponentially, often surpassing the initial damage inflicted by the windstorm itself. This pattern is evident in both study areas (Figure 7); according to the Forwind dataset, which tracks also the areas affected by Vaia windstorm, and the freely available data of bark beetle infestation shared by the local forest services, the bark beetle outbreak over two years has caused damage nearly equivalent (for the Belluno area) or even higher (for Bolzano area) to that caused by the Vaia windstorm. This exponential increase in damage highlights the cascading effects of compound disturbances, as discussed in studies on forest resilience and disturbance interactions [5, 80].

Our research represents the first attempt to analyze this correlation for the Vaia windstorm, incorporating additional factors such as structural forest conditions and meteorological variables that might influence the relationship. Our analysis indicates that, the damage caused by the Vaia windstorm may have been a contributing factor to the bark beetle outbreak. On the other side, our results suggest that almost all the forest structural variables (except the forest HH in the Belluno area), estimated using LiDAR data, are similar in all areas affected by the bark beetle, whether they intersect the regions impacted by the Vaia windstorm or in other *Picea abies* forests not affected by the windstorm, indicating that the considered forest structure variables, as assessed through our LiDAR approach, may not be a trigger for the outbreak. Recent studies showed that other structural variables such as crown closure, stand age, and elevation have been consistently identified as influential factors in predicting bark beetle (*Ips sexdentatus*) susceptibility [86, 59]. While our findings using LiDAR-derived metrics did not strongly correlate with outbreak risks in the Vaia context, these studies that focused the infestation of *Ips sexdentatus* in crimean pine forests underscores the critical role of stand composition and structure. For instance, Sivrikaya et al. [86] demonstrated the utility of combining analytical hierarchy process, frequency ratio, and logistic regression models to map these infestations susceptibility emphasizing crown closure and elevation as predominant factors, aligning with the outcomes of Özcan et al. [59], who highlighted the significant roles of maximum temperature and solar radiation alongside forest structure.

Researches has shown that drought conditions can significantly favor bark beetle outbreaks by stressing trees, making them more vulnerable to infestation [56, 49, 55]. Studies indicate that drought weakens the physiological resistance of trees like *Picea abies*, creating ideal conditions for bark beetles to thrive and proliferate. In both our study areas, however, the results showed that areas affected by Vaia windstorm and subsequent bark beetle infestations had similar temperatures compared to other *Picea abies* forests with bark beetle presence but not impacted by Vaia. Precipitation exhibited contrasting patterns: in the Bolzano area, higher precipitation values were observed in regions affected by the Vaia windstorm and subsequent bark beetle infestations, whereas in the Belluno area, higher precipitation was found in other *Picea abies* forests with bark beetle presence. This difference can be attributed to the specific areas hit by Vaia windstorm, which are predominantly mountainous regions (in the northern part of the Belluno Province and in the south and east part of the Bolzano Province) characterized localized precipitation than the rest of the Provinces.

When we considered all the meteorological and structural variables together in both study areas, we found that none of them were ultimately significant for predicting the interaction between bark beetle infestations and areas affected by the Vaia windstorm. This further emphasizes that the considered variables may not be triggers for the outbreak.

Our analysis indirectly considered other factors such as species composition and altitude. We focused our study exclusively on spruce forests, the most common species in the area (the most hit by the Vaia windstorm) and the primary species affected by the bark beetle [47]. Additionally, we accounted for altitude by stratifying our data randomly based on altitude gradients. This comprehensive approach ensures that our findings are robust and account for the key variables that could influence the correlation between the Vaia windstorm and the bark beetle outbreak.

Other factors might have significantly influenced the observed correlation. As stressed in different studies, forest management is essential in mitigating the impacts of such compounded events [56, 47] by employing multiple strategies to enhance forest resilience and resistance [38]. As climate change continues to increase the frequency and severity of these disturbances, we must anticipate and prepare for more frequent occurrences of such events [79]. Preventive strategies aim at maintaining low beetle population densities [84, 35], focusing on creating robust forests capable of withstanding various disturbances, including large-scale stochastic mortality events and the complex ecological effects of phenomena such as windstorms. Implementing silvicultural practices that promote heterogeneity and mixed stand structures can further enhance forest resilience and reduce vulnerability to windstorm damage than less diverse stands [38, 34] consequently reducing the risk of beetle damage in Norway spruce. Another preventive measure is to establish a comprehensive network of forest roads and support points to facilitate more effective active forest management. However, in different areas such as in mountainous terrain (as in our study areas), both detailed forest management and the construction of new roads are particularly challenging due to the rugged topography.

After wind damage, forest management should focus on salvage logging, aiming to collect all fallen trees as quickly as possible, which can accumulate becoming fuel and a possible risk for forest fires [46]. However, in largescale events like the Vaia windstorm, which affected thousands of hectares and uprooted numerous trees, it is challenging to promptly collect all the fallen timber through salvage logging. Consequently, a subsequent bark beetle outbreak is likely to occur. Additionally, in mountainous regions, such as in our study, intervention is particularly challenging due to the rugged topography, difficult access to many areas, and risks such as falling rocks or snow. However, the question remains open as to whether salvage logging in areas impacted by massive and intense windstorms is effective in preventing future disturbances from bark beetles [43]. Instead, salvage logging might potentially increase the risk of further wind damage, thereby offsetting the benefit of reduced bark beetle disturbances [17].

Continued research and adaptive management practices are essential to safeguard the forest ecosystems in the face of a changing climate. Fostering forest resilience and adaptive capacity will be vital in mitigating the adverse effects of climate change on forest health and productivity [79]. Further research is needed to explore these additional variables and to better understand the complex interplay of factors contributing to bark beetle outbreaks.

It is worth underlying that this research was made possible by the availability of open-source data, including the Forwind dataset (for the estimation of forest hit by Vaia windstorm), the bark beetle infestation dataset (from both local forest services) and the meteorological data. Also the importance of remote sensing information (particularly LiDAR) in enhancing our understanding of forest structure and dynamics, was clear in this study. LiDAR’s ability to capture detailed three-dimensional information efficiently and cost-effectively, along with its capability to cover extensive areas quickly, positions it as an indispensable tool for ecosystem monitoring [29]. While there may be trade-offs in accuracy compared to traditional field data collection methods, LiDAR provides a practical and expedient alternative for assessing large-scale disturbances [33, 1]. The methodology employed to estimate structural variables in our study utilized a per-pixel approach, which, unlike the per-point cloud approach [93], involves raster analysis of the CHM. This method provides a broad overview of forest structure across large areas efficiently and economically. It excels in rapidly capturing extensive data, although it may generalize individual tree characteristics, potentially affecting the accuracy of forest density and HH estimates. Another factor of uncertainties could be related to the spatial resolution of the CHM used to calculate the aforementioned structural variables that, as as previously stated differed between the study areas: 1m for Belluno (collected in 2015) and 2.5m for Bolzano (collected in 2006). We believe that this difference in CHM spatial resolution and the respective LiDAR acquisition dates does not significantly impact our research results. As demonstrated by Torresani et al. [96], small differences in spatial resolutions and relatively minor time mismatches between data acquisition and analysis yield similar results for HH estimation in forestry areas. This is a common issue when working with ALS LiDAR data across different regions, as not all areas have uniform ALS LiDAR coverage.

The choice of the heterogeneity index for the estimation of HH, can be considered another point of uncertainties of this study. For HH estimation, we decided to rely on the Rao’s Q index, a metric used in differnet studies [87, 96, 94], including those focusing on the Spectral Variation Hypothesis [97, 72, 89, 61] that showed robust results. Rao’s Q index integrates both relative abundance and pixel values through Euclidean distance, comprehensively captures the structural information derived from LiDAR data heterogeneity [92, 62]. When applied to a single layer or raster, such as the CHM as in our study, Rao’s Q serves as a robust proxy for heterogeneity, converging to variance using half the squared Euclidean distance (1/2 *d_ij_*^2^). The decision to focus on the use of CHM and not to other variables, is supported by Tamburlin et al. [87], who conducted a thorough exploration of various LiDAR metrics (such as entropy, standard deviation of point cloud distribution, and percentage of returns above mean height) for HH estimation, their findings indicated that the CHM was the most effective metric for characterizing HH. Wrapping up the discussion on the data used in this study, it is important to emphasize the significance of openly accessible datasets (such as the Forwind dataset, ALS LiDAR data, and bark beetle infestation data) and algorithm functions (like the *rasterdiv* R package) for ecological research. These resources are vital, allowing scientists to perform comprehensive analyses and derive meaningful conclusions about environmental phenomena [73]. The availability of these comprehensive datasets, allows for a more thorough understanding of complex ecological interactions and the development of informed management strategies. Our research demonstrates how integrating open-source data with advanced remote sensing technologies like LiDAR can provide invaluable insights into forest dynamics and disturbances. This approach not only enhances the accuracy and efficiency of ecological monitoring but also underscores the importance of maintaining and expanding open-access data repositories to support ongoing and future research efforts in environmental science.

## 5 Conclusion

The findings of this study highlight the possible relationship between windstorm events, such as the Vaia windstorm, and subsequent bark beetle outbreaks. Notably, the study indicates that structural variables estimated through LiDAR data, do not play a significant role in this correlation. It remains to be tested whether a broader spectrum of stand structural traits, determined by various forest management practices and resulting in a less homogeneous forest landscape, would lead to different correlation outcomes. Nevertheless, the approach utilized in the present study is undoubtedly useful for investigating the effects of alternative disturbance management strategies. As climate change continues to increase the frequency and severity of these disturbances, we must anticipate and prepare for more frequent occurrences of such events [79]. This underscores the necessity of developing and adopting an adaptable management framework, such as Climate Smart Forestry [90], to enhance the resilience and sustainability of forest ecosystems, characterized by higher tree species diversity and complex forest structures [38, 79]. In this context, advances in remote sensing tools, which offer a detailed view of forest structure and composition, are essential for understanding the environmental mechanisms behind the variability in disturbance impacts across complex mountain landscapes.

## 6 Author contributions

MT: Conceptualization, Data curation, Formal analysis, Investigation, Methodology, Visualization, Writing – original draft. RT: Conceptualization, Formal analysis, Investigation, Methodology, Funding acquisition, Project administration, Supervision, Validation, Writing – original draft.

